# Sub-type selective muscarinic acetylcholine receptors modulation for the treatment of parkinsonian tremor

**DOI:** 10.1101/2022.04.04.487007

**Authors:** Kinsey Bickham, C. Price Withers, Augusto Diedrich, Mark Stephen Moehle

## Abstract

Parkinson’s Disease is characterized by hallmark motor symptoms including resting tremor, akinesia, rigidity, and postural instability. In patient surveys of Parkinson’s Disease symptoms and quality of life, tremor consistently ranks among the top concerns of patients with disease. However, the gold standard of treatment, levodopa, has inconsistent or incomplete anti-tremor effects in patients, necessitating new therapeutic strategies to help relieve this burden. Non-selective anti-muscarinic acetylcholine receptor therapeutic agents which target each of the 5 muscarinic receptor subtypes have been used as an adjunct therapy in this disease, as well as other movement disorders, and have been shown to have anti-tremor efficacy. Despite this, anti-muscarinic therapy is poorly tolerated due to adverse effects. Recent pharmacological advances have led to the discovery of muscarinic subtype selective antagonists that may keep the anti-tremor efficacy of non-selective compounds, while reducing or eliminating adverse effects. Here, we directly test this hypothesis using pharmacological models of parkinsonian tremor combined with recently discovered selective positive allosteric modulators and antagonists of the predominant brain expressed muscarinic receptors M_1_, M_4_, and M_5_. Surprisingly, we find that selective modulation of M_1_, M_4_, or M_5_ does not reduce tremor in these pre-clinical models, suggesting that central or peripheral M_2_ or M_3_ receptors may be responsible for the anti-tremor efficacy of non-selective anti-muscarinic therapies currently used in the clinic.

## Introduction

Parkinson’s disease (PD) is the second most common neurodegenerative disorder worldwide, with a prevalence of 1% for individuals above 60 years of age(1). PD is characterized by resting tremor, rigidity, akinesia/bradykinesia, and postural instability, as well as several non-motor symptoms(2). The current standard of treatment for PD is dopamine (DA) replacement through the administration of the DA precursor levodopa (L-dopa). However, chronic L-dopa treatment can lead to severe adverse effects or is not highly efficacious at treating certain PD motor symptoms, such as resting tremor and gait and several non-motor symptoms(1, 3).

Resting tremor is consistently reported as the one of the most troublesome motor symptoms of PD(4-6). In PD patient surveys, tremor is among the most frequently mentioned symptoms, and it is consistently ranked as the most important symptom to alleviate(5). Additionally, tremor is one of the most common symptoms that drives neurologists to modify ongoing anti-parkinsonian drug treatments(4). Unfortunately, tremor response to L-dopa is highly variable(6), highlighting the unmet clinical need of new therapeutic options targeting tremor in PD patients.

Anti-muscarinic therapeutics which target each of the 5 muscarinic acetylcholine receptor subtypes (mAChRs, M_1_-M_5_) were among the first widely accepted treatments for PD (7, 8). Anti-mAChRs have efficacy in reducing parkinsonian tremor(7). Despite their efficacy, non-selective anti-mAChR therapy has limited clinical utility due to the potentially severe peripheral and central adverse effects(3, 9, 10). Recent advances in our understanding of the roles of each mAChR subtype have yielded the possibility that targeting individual mAChRs may maintain the efficacy observed with non-selective therapeutics while reducing or eliminating the adverse effects(10, 11). Yet, this requires further knowledge of each receptor’s activity in both pre-clinical models of disease as well as normal physiological processes.

Preclinical studies suggest that M_2_ and M_3_ underly the adverse peripheral side effects associated with non-selective anti-mAChR therapeutics, and, while expressed in the central nervous system, have higher expression in peripheral organs(3, 11-14). Conversely, M_1_, M_4_, and M_5_ are the brain predominant mAChRs and have higher expression in the central nervous system compared to the periphery(11, 12). M_1_, M_4_, and M_5_ are expressed in multiple brain regions associated with PD pathophysiology including throughout several basal ganglia nuclei, cortical regions, and substantia nigra pars compacta dopaminergic cells (12, 15). These expression patterns suggest that the anti-parkinsonian efficacy of non-selective compounds may be through inhibition of M_1_, M_4_, and M_5_. Initial studies in 6-OHDA lesioned mice have suggested that this may indeed be the case with antagonism of M_1_ and M_4_ displaying anti-parkinsonian efficacy (16, 17). However, many of these studies have largely relied on non- or poorly-selective mAChR tool compounds and have not directly assessed parkinsonian tremor.

Parkinsonian tremor can be modeled in rodents with tremulous jaw movements (TJM)(18). These are vertical deflections of the rodent jaw that occur in a similar 3-7.5 Hz frequency range to the tremor of patients with PD (18, 19).TJMs are induced pharmacologically with dopamine depleting agents such as tetrabenazine and pro-cholinergic agents such as acetylcholinesterase inhibitors. TJMs can be reduced in frequency or severity by agents that treat tremor in PD patients including pro-dopaminergic therapies and non-selective anti-mAChRs (18, 19), suggesting that this model has face and construct validity in determining novel therapeutics that will display anti-tremor efficacy in PD patients.

In this study, we used recently developed M_1_, M_4_, and M_5_ mAChR subtype selective antagonists and positive allosteric modulators to manipulate the activity of M_1_, M_4_, and M_5_ *in vivo* in relevant rodent models of parkinsonian tremor to understand how individual mAChRs regulate tremor. To accomplish this, we utilized two pharmacological agents, galantamine and tetrabenazine, which have distinct mechanisms of action to induce TJMs in mice. Galantamine works through increasing levels of acetylcholine by inhibiting this neurotransmitter’s breakdown(20). Tetrabenazine induces TJMs through depletion of monoamines(19). Notably, both of the agents have been extensively used to induce TJMs previously. Using these highly selective compounds, as well as a non-selective anti-mAChR antagonist as a positive control, in this pharmacologically-induced TJM model we systematically tested the role of the brain predominant mAChRs in modulating tremor.

## Materials and Methods

### Animals

C57Bl6/J mice were purchased from Jackson Labs (Stock #000664). Male mice were purchased between 8-10 weeks of age, and all mice were used at ages starting at 10-12 weeks of age. Mice were maintained in AALAS approved vivariums on 12 hour light/dark cycles with *ad libitum* access to food and water. All studies were approved by the institutional animal care and use committee.

### Pharmacological Agents

VU0467154(21), VU06021625(9), VU0453595 (22), VU0255035 (23), and ML375 (24) were synthesized by the medicinal chemists of the Warren Center for Neuroscience Drug Discovery according to published protocols. Tetrabenazine, galantamine, and scopolamine were purchased from Tocris. VU0467154, VU0453595, and VU0255035 were dissolved in 10% Tween-80 (w/v) in sterile saline. VU0601625 and ML375 were dissolved in 20% hydroxy-propyl-β-cyclodextrin. Galantamine and scopolamine were dissolved in sterile saline. Tetrabenazine was dissolved in 10% DMSO (w/v) in sterile water that had been acidified with 0.1% 1M HCL, and then pH adjusted once dissolved. All drugs were made fresh daily.

### Tremulous Jaw Movements

Mice were injected intraperitoneally with tetrabenazine or galantamine – 120 minutes and 30 minutes before recording, respectively. Each mAChR test ligand was injected subcutaneously 30 minutes in advance of recording. Ten minutes before recording, mice were placed on the observational setup for a habituation period; recordings lasted for 15 minutes. The observational chamber was a clear Plexiglas tube 10 cm in diameter with a mesh floor. The video camera recorded a ventral view of the mice through the mesh floor. Mice received each dose of drug in each drug group plus a vehicle and scopolamine in a within-subject design. With the exception of the dose responses curves of tetrabenazine and galantamine which were scored live, all videos were scored at a later date by reviewers blinded to the experimental conditions.

### Data Analysis

Videos were reviewed using Final Cut Pro by experimenters trained in the analysis of tremulous jaw movements. This program allows for frame-by-frame analysis of movement. Experimenters were blinded to drug treatment and dose. All statistical analyses were performed in GraphPad Prism 9 (GraphPad, San Diego, CA). Statistical tests were used throughout the paper with the following scheme: If the data were normally distributed as determined by a D’Agonstino and Pearson omnibus normality test, a repeated measures ANOVA was utilized with either a Sidak’s or Dunnett’s post-hoc comparison. If the data were not normally distributed a Friedman’s test with Dunnett’s post comparison was used. Because of the within subjects design, if data were missing, a Mixed Effects Model was used to analyze the data.

## Results

### M4 Antagonists Do Not Reduce TJMs

Before testing our mAChR test compounds, we first empirically determined the doses of tetrabenazine and galantamine that produced maximal TJMs and a threshold amount of TJMs. We administered 1, 3, or 10 mg/kg (intraperitoneally (i.p.)) of tetrabenazine or 0.3, 1, 2 or 3 mg/kg of galantamine i.p to C57Bl6/J mice and observed TJMs. For tetrabenazine, 3 and 10 mg/kg i.p significantly induced TJMs over vehicle treatment with 10 mg/kg producing a maximal effect and 3 mg/kg producing a threshold effect (Figure S1 A, One-way ANOVA with Dunnet’s multiple comparisons test F(4,67)=31.72 p<0.0001, p<0.01 for 3 mg/kg, p<0.0001 for 10 mg/kg). For galantamine, 1 and 3 mg/kg i.p significantly raised induced TJMs over vehicle treatment with 3 mg/kg producing a maximal effect and 1 mg/kg producing a threshold effect (Figure S1 B, One-way ANOVA with Dunnet’s multiple comparisons test F(3,36)=44.92 p<0.0001, p<0.05 for 1 mg/kg, p<0.0001 for 3 mg/kg).

Recent evidence suggests a unique role for M_4_ antagonists in the treatment of movement disorders(3, 9). To determine if M_4_ selective antagonists can recapitulate the effects of non-selective antagonists we performed a dose response curve of the M_4_ selective antagonist VU06021625 (0, 3, 10 or 30 mg/kg s.c.) in the presence of maximally efficacious doses of tetrabenazine (10 mg/kg i.p, Figure 1A) or galantamine (3 mg/kg i.p, Figure 1B). We additionally dosed the animals with scopolamine (1 mg/kg, s.c.) in the presence of these TJM-inducing pharmacological agents as a positive control. In tetrabenazine induced TJMs, VU6021625 did not significantly alter TJM counts (Figure 1A, Mixed Effects Model with Holm-Sidak multiple comparisons test, p>0.05). However, scopolamine significantly attenuated TJMs (Figure 1A, Mixed Effects Model with Holm-Sidak multiple comparisons test, p<0.05). For galantamine induced TJMs, VU06021625 surprisingly facilitated tremor at higher doses while scopolamine significantly attenuated tremor (Figure 1B, Repeated measures one-way ANOVA with Sidak’s multiple comparisons test F(2.792,25.013)=18.26 p<0.0001, p<0.05 for 10 mg/kg, p<0.05 for 30 mg/kg p<0.0001 for scopolamine). Taken together, this suggests that M_4_ selective antagonists do not recapitulate the effects of non-selective antagonists and may even potentiate tremor in some circumstances.

**Figure 1.**
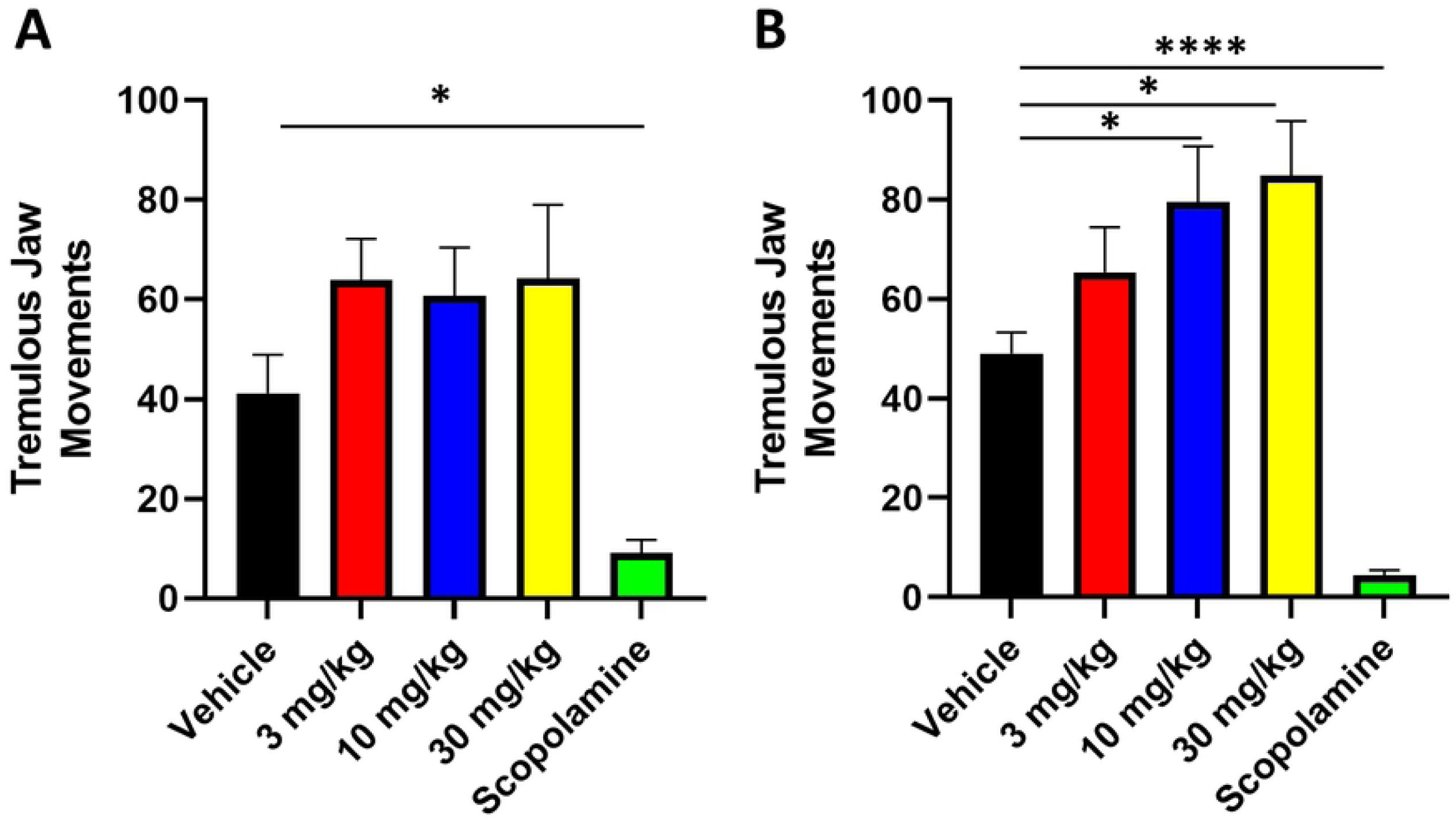
M_4_ Selective Antagonists Do Not Recapitulate Non-selective Antagonists. **A)** Dose response curve of VU06021625 (0-30 mg/kg s.c.) in tetrabenazine induced tremulous jaw movements. The positive control scopolamine (1 mg/kg s.c.) was the only compound that reduced tremor. **B)** Dose response curve of VU06021625 (0-30 mg/kg s.c.) in galantamine induced tremulous jaw movements. Higher doses of VU06021625 potentiated TJM counts while scopolamine (1 mg/kg s.c.) decreased TJM counts. N=10 per group. Data in A were analyzed with a mixed effects model with Holm-Sidak multiple comparisons test. Data in B were analyzed with a repeated measures one-way ANOVA with Sidak’s multiple comparisons test. *p<0.05, and ****p<0.0001

### M1 Antagonists Do Not Reduce TJMs

M_1_ has previously been implicated in having a role in modulating the parkinsonian basal ganglia(16, 17). To determine if M_1_ selective antagonists can recapitulate the effects of non-selective antagonists we performed a dose response curve of the M_1_ selective antagonist VU0255035 (0, 1, 3, or 10 mg/kg s.c.) in the presence of maximally efficacious doses of tetrabenazine (10 mg/kg i.p, Figure 2A) or galantamine (3 mg/kg i.p, Figure 2B). As before, we additionally dosed the animals with scopolamine (1 mg/kg, s.c.) in the presence of these TJM inducing pharmacological agents as a positive control. In tetrabenazine or galantamine induced TJMs, VU0255035 did not significantly alter TJM counts (Figure 2A (tetrabenazine), Repeated measures one-way ANOVA with Dunnett’s multiple comparisons test F(2.559, 17.91)=4.428 p<0.05, not significant for all doses, Figure 2B (galantamine) Repeated measures one-way ANOVA with Dunnett’s multiple comparisons test F(1.828, 12.79)=5.5357, p<0.05, not significant for all doses). However, scopolamine significantly attenuated TJMs in both models (Figure 2A (tetrabenazine), Repeated measures one-way ANOVA with Dunnett’s multiple comparisons test F(2.559, 17.91)=4.428 p<0.05, p<0.01 for scopolamine, Figure 2B (galantamine) Repeated measures one-way ANOVA with Dunnett’s multiple comparisons test F(1.828, 12.79)=5.5357, p<0.05, p<0.001 for scopolamine). This suggests that M_1_ selective antagonists do not recapitulate the effects of non-selective antagonists.

**Figure 2.**
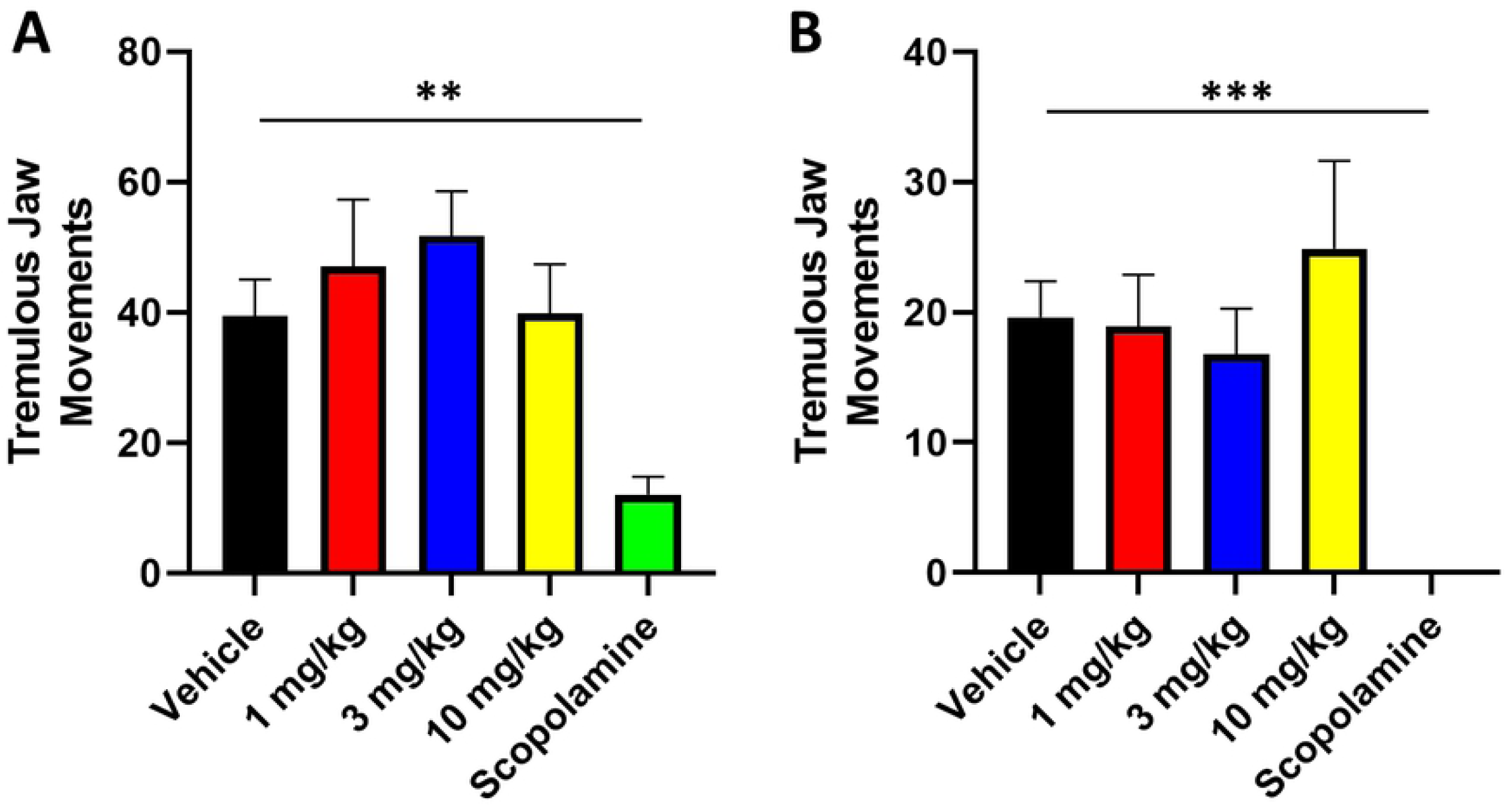
M_1_ Selective Antagonists Do Not Recapitulate Non-selective Antagonists. **A)** Dose response curve of VU0255035 (1-10 mg/kg s.c.) in tetrabenazine induced tremulous jaw movements. The positive control scopolamine (1 mg/kg s.c.) was the only compound that reduced tremor. **B)** Dose response curve of VU0255035 (1-10 mg/kg s.c.) in galantamine induced tremulous jaw movements. Only Scopolamine significantly reduced TJM counts N=8-9 per group. Data were analyzed with a repeated measures one-way ANOVA with Dunnett’s multiple comparisons test. **p<0.01, and ***p<0.001

### M5 Antagonists Do Not Reduce TJMs

M_5_ is unique among mAChRs in that its expression is largely limited to midbrain dopaminergic neurons(15, 25, 26). To determine if M_5_ selective antagonists can recapitulate the effects of non-selective antagonists we performed a dose response curve of the M_5_ selective negative allosteric modulator ML375 (0, 3, 10, or 30 mg/kg s.c.) in the presence of maximally efficacious doses of tetrabenazine (10 mg/kg i.p, Figure 3A) or galantamine (3 mg/kg i.p, Figure 3B). In tetrabenazine or galantamine induced TJMs, ML375 did not significantly alter TJM counts (Figure 3A (tetrabenazine), Friedman’s test with Dunnett’s multiple comparisons test Friedman=16.00 p<0.05, not significant for all doses, Figure 2B (galantamine) Repeated measures one-way ANOVA with Dunnett’s multiple comparisons test F(2.452, 17.16)=1.217, p<0.05, not significant for all doses). However, scopolamine significantly attenuated TJMs in the TBZ model (Figure 3A Friedman’s test with Dunnett’s multiple comparisons test Friedman=16.00 p<0.05, p<0.05 for scopolamine). Interestingly, when all vehicle and scopolamine data were pooled from all antagonist groups, scopolamine was always highly efficacious in reducing TJM counts in either pharmacological model (Figure S2. Paired t test (A) t=8.350 df=23, (B) t=8.292 df=18, p<0.0001). This suggests that M_5_ selective antagonists do not recapitulate the effects of non-selective antagonists.

**Figure 3.**
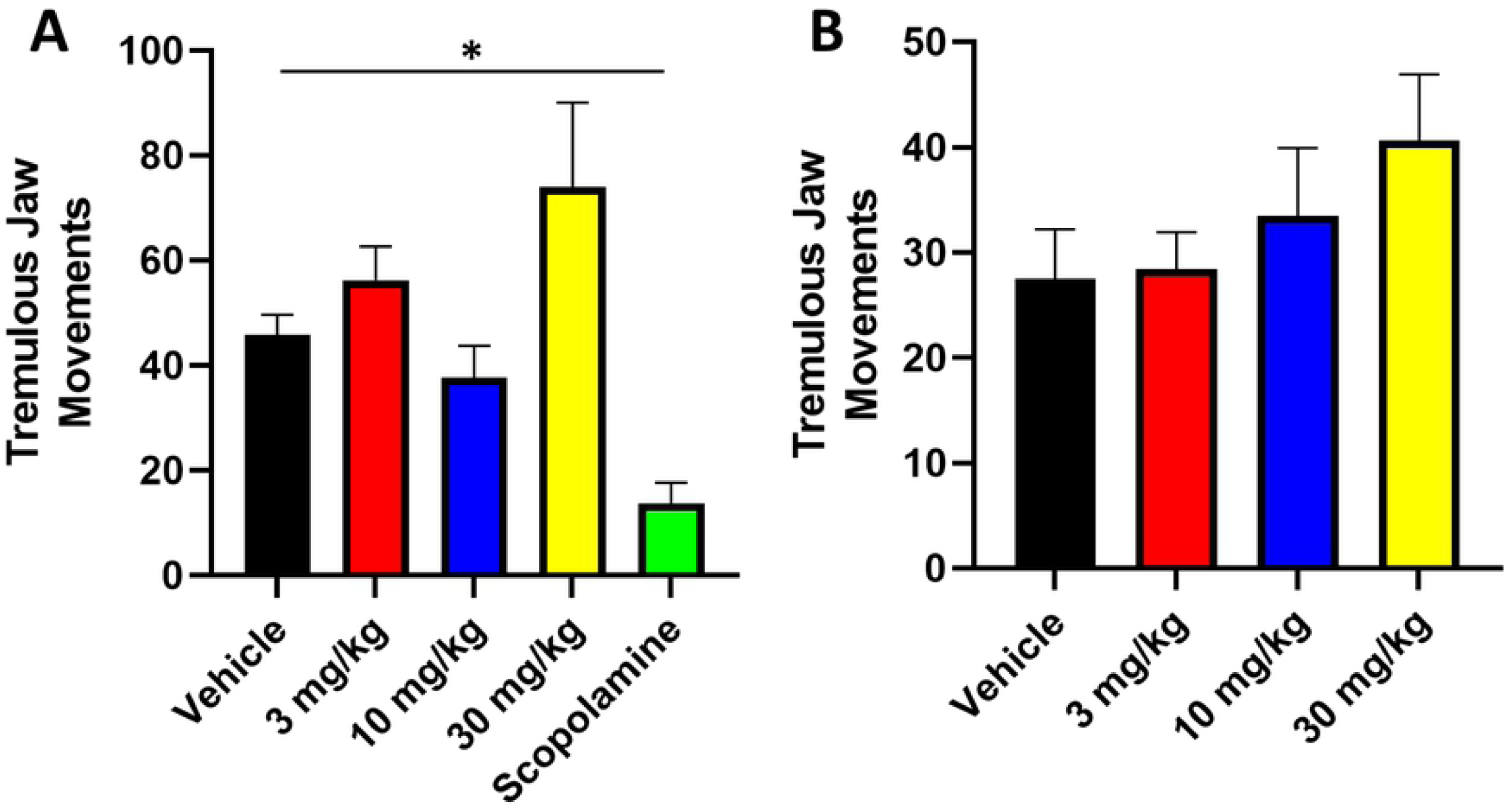
M_5_ Selective Antagonists Do Not Recapitulate Non-selective Antagonists. **A)** Dose response curve of ML375 (3-30 mg/kg s.c.) in tetrabenazine induced tremulous jaw movements. The positive control scopolamine (1 mg/kg s.c.) was the only compound that reduced tremor. **B)** Dose response curve of ML375 (3-30 mg/kg s.c.) in galantamine induced tremulous jaw movements. N=7-8 per group. Data in A were analyzed with a Friedman’s test with a Dunnett’s multiple comparisons test. Data in B were analyzed with a repeated measures one-way ANOVA with Dunnett’s multiple comparisons test. *p<0.05

### M4 PAMs Do Not Modulate TJMs

Due to our results suggesting that M_4_ antagonism does not decrease TJMs, we tested if the reverse experiment, boosting M_4_ signaling, would alleviate tremor in a manner similar to non-selective mAChR antagonists. To determine if M_4_ selective positive allosteric modulators can alleviate tremor we performed a dose response curve of the M_4_ selective positive allosteric modulator VU0467154 (0, 0.3, 1, or 3 mg/kg s.c.) in the presence of a minimally efficacious doses of tetrabenazine (3 mg/kg i.p, Figure 4A) or galantamine (1 mg/kg i.p, Figure 4B). In tetrabenazine or galantamine induced TJMs, VU0467154 did not significantly increase TJM counts (Figure 4A (tetrabenazine), Repeated measures one-way ANOVA with Dunnett’s multiple comparisons test F(2.6, 15.60)=5.038 p<0.05, not significant for all doses, Figure 2B (galantamine) Repeated measures one-way ANOVA with Dunnett’s multiple comparisons test F(2.980, 23.84)=6.536, p<0.05, not significant for all doses). This suggests increasing M_4_ activity through M_4_ selective positive allosteric modulators do not modulate TJMs.

**Figure 4.**
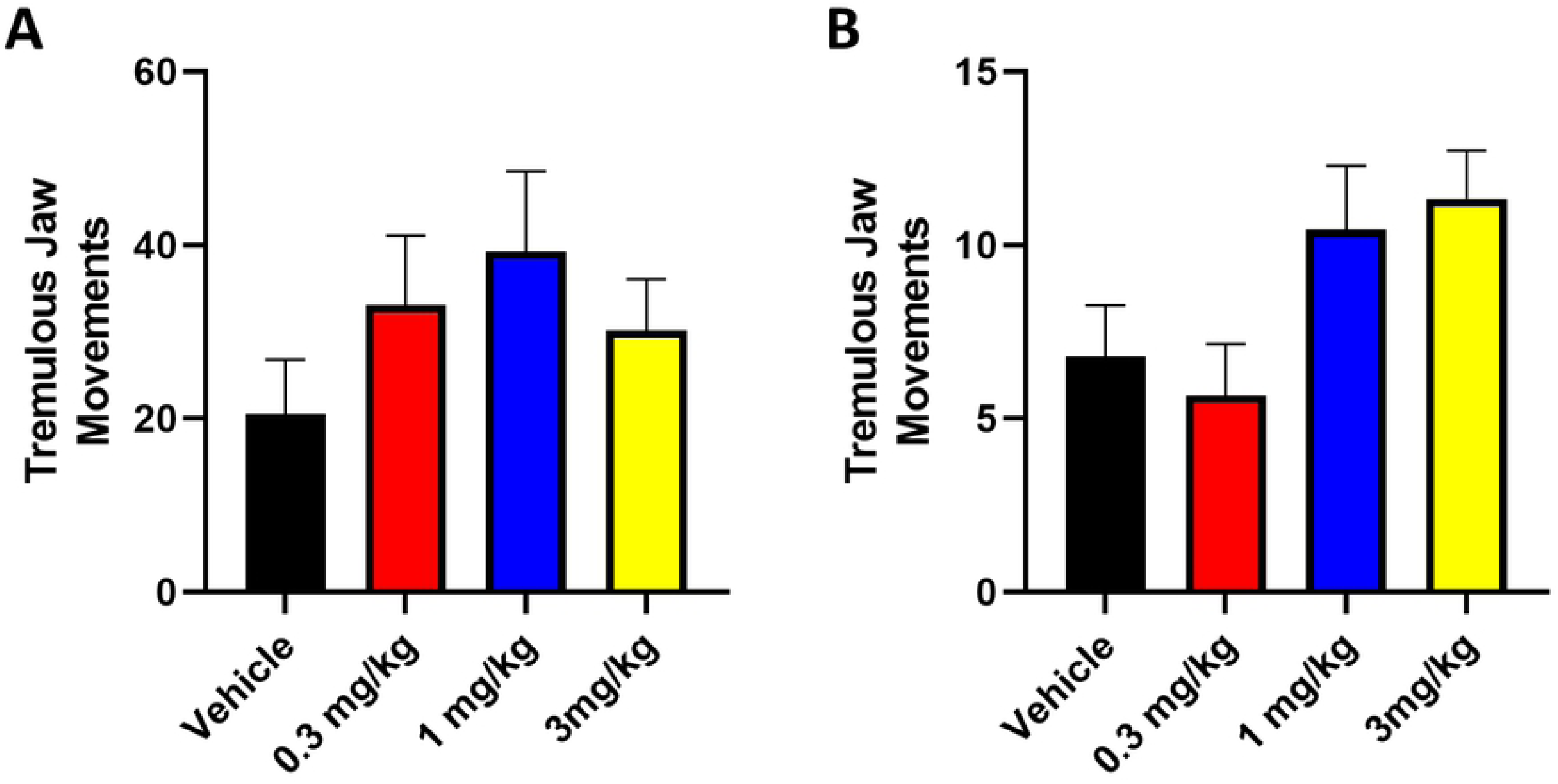
M_4_ Selective Positive Allosteric Modulators Do Not Modulate Tremulous Jaw Movements. **A)** Dose response curve of VU0467154 (0.3-3 mg/kg s.c.) in modulating a threshold dose of tetrabenazine. **B)** Dose response curve of VU0467154 (0.3-3 mg/kg s.c.) in modulating a threshold dose of galantamine induced tremulous jaw movements. N=7-9 per group. Data were analyzed with a repeated measures one-way ANOVA with Dunnett’s multiple comparisons test

### M1 PAMs Do Not Modulate TJMs

As with M_4_ modulation, we tested if the reverse experiment, boosting M_1_ signaling, would alleviate tremor in a manner similar to non-selective mAChR antagonists. To determine if M_1_ selective positive allosteric modulators can alleviate tremor we performed a dose response curve of the M_1_ selective positive allosteric modulator VU0453595 (0, 1, 3, or 10 mg/kg s.c.) in the presence of a minimally efficacious doses of tetrabenazine (3 mg/kg i.p, Figure 4A) or galantamine (1 mg/kg i.p, Figure 4B). In tetrabenazine or galantamine induced TJMs, VU0453595 did not significantly increase TJM counts (Figure 5A (tetrabenazine), Repeated measures one-way ANOVA with Dunnett’s multiple comparisons test F(1.607, 11.25)=2.703 p>0.05, not significant for all doses, Figure 5B (galantamine), Mixed Effects Model with Holm-Sidak multiple comparisons test, p>0.05, not significant for all doses). This suggests increasing M_1_ activity through M_1_ selective positive allosteric modulators do not modulate TJMs.

**Figure 5.**
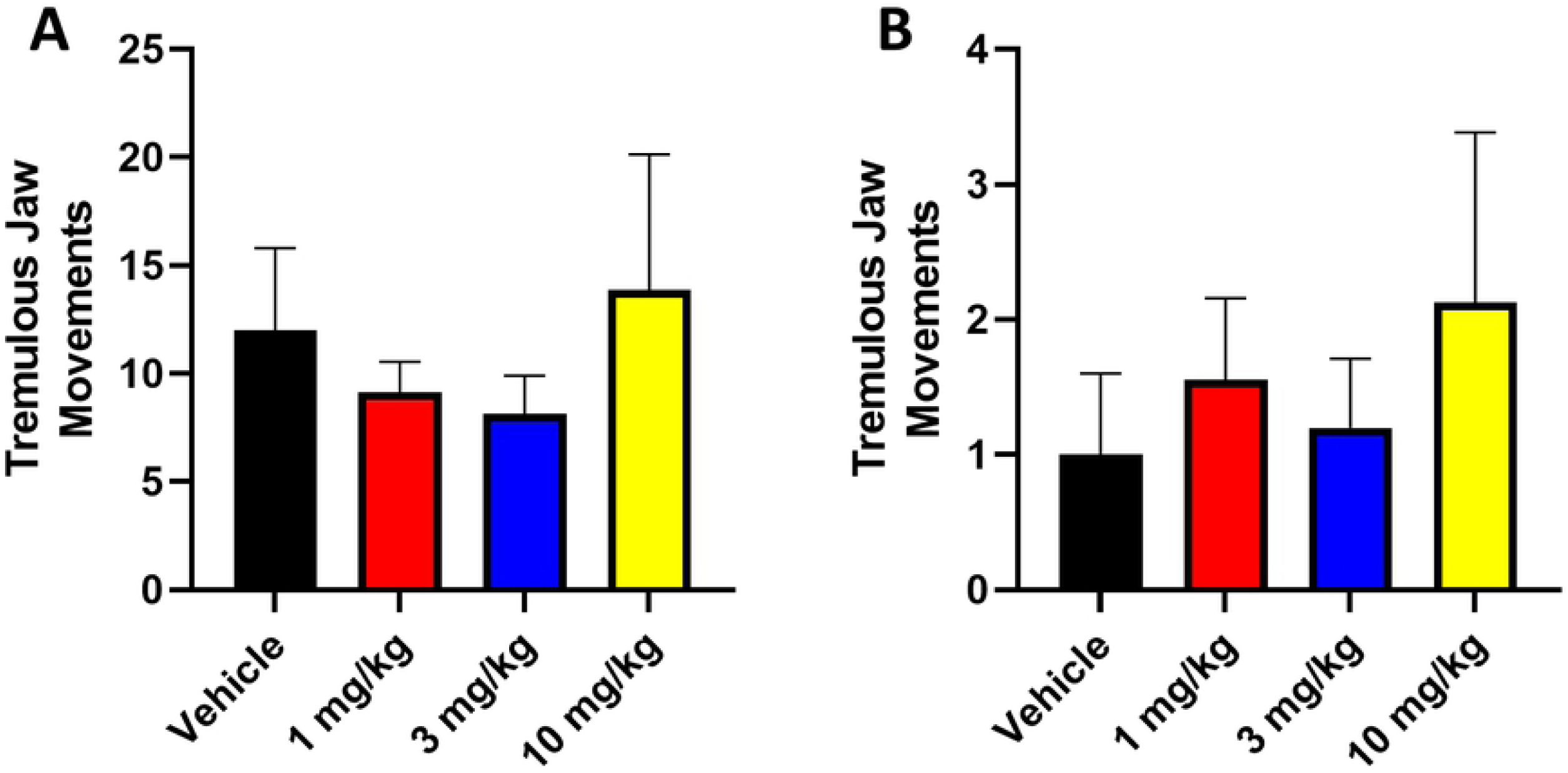
M_1_ Selective Positive Allosteric Modulators Do Not Modulate Tremulous Jaw Movements. **A)** Dose response curve of VU0453595 (1-10 mg/kg s.c.) in modulating a threshold dose of tetrabenazine. **B)** Dose response curve of VU0453595 (1-10 mg/kg s.c.) in modulating a threshold dose of galantamine induced tremulous jaw movements. N=8-10 per group. Data in A were analyzed with a repeated measures one-way ANOVA with Dunnett’s multiple comparisons test. Data in B were analyzed with a mixed effects model with Holm-Sidak multiple comparisons test

## Discussion

Pharmacological advances have allowed for the development of truly mAChR subtype selective pharmacological tool compounds that enable dissection of the roles of individual mAChR subtypes in normal physiology and in disease states(3, 10, 11). These advances have placed a renewed excitement in utilizing cholinergic agents clinically as the adverse effects traditionally associated with mAChRs may be avoided or greatly reduced. However, careful pre-clinical work must be done to understand the extent and mechanisms of potential efficacy of these selective compounds.

Recent studies have implicated that, in pre-clinical models of anti-parkinsonian efficacy, antagonism of M_4_ underlies the majority of efficacy seen with non-selective mAChR antagonists(9). Previous studies have also implicated a role for M_1_ (17). However, these animal models, forelimb asymmetry and haloperidol induced catalepsy, primarily model the hypokinetic motor deficits of PD while tremor is hyperkinetic. They implicate a unique role for M_1_ and M_4_ in the physiology of the hypokinetic motor symptoms of PD. Our data using highly selective pharmacological tools to both antagonize or potentiate M_1_, M_4_, and M_5_ implicate that these brain predominant mAChRs are not the primary mAChRs that are responsible for modulating tremor. This indicates a possible dissociation of the mechanisms or circuits which underlie hypokinetic and hyperkinetic motor symptoms of PD.

The only departure to a lack of efficacy in our data is M_4_ antagonists potentiating tremor in the galantamine model. It should be noted that this only happens at doses in excess of those necessary for anti-parkinsonian and anti-dystonic efficacy, suggesting that M_4_ antagonists will be able to relieve hypokinetic motor deficits without potentiation other motor symptoms(9). The differences between effects in the tetrabenazine and galantamine TJM model may also indicate that distinct mechanisms are engaged to induce tremor. For example, based on the expression profile of M_4_ in the striatum, it may be possible that M_4_ antagonism increases TJM through inhibiting M_4_ on striatal cholinergic interneurons and further increased acetylcholine concentrations in the striatum to induce tremor, whereas this cholinergic interneuron mechanism may not be engaged by tetrabenazine administration(18-20, 27).

The lack of efficacy seen with M_1_, M_4_, and M_5_ antagonists in our pharmacological models of TJMs indicates that the observed efficacy of non-selective antagonists may be due to antagonism of M_2_ or M_3_ mAChRs. Previous studies have also suggested the possible role of M2 antagonism in tremor, as global M_2_ knockout mice were less sensitive to the tremor inducing mAChR agonist oxotremorine(28). Due to the classic peripheral adverse effects associated with non-selective mAChR therapeutics possibly being linked to actions at M_2_ or M_3_, this may indicate that targeting mAChRs for tremor may not be dissociable from adverse effects, and other novel strategies for decreasing tremor in PD may be warranted.

Our data also raise the question of which circuits and organs anti-mAChR therapeutics are acting on for anti-tremor efficacy. While not as highly expressed as M_1_, M_4_, and M_5_ are in the brain, nevertheless, M_2_ and M_3_ are still found in the brain. Notably, M_2_ is expressed on cholinergic interneurons in the striatum, which are critical regulators of basal ganglia processing(12, 15). Additionally, M_3_ is expressed in several brain nuclei which may influence basal ganglia processing associated with PD pathophysiology(12, 15). This expression profile suggests that there may be several central mechanisms by which antagonism of M_2_ or M_3_ may alter tremor.

M_2_ and M_3_ are also expressed throughout several peripheral sites including muscle and lower motor neurons(15). Our data cannot rule out the possibility that non-selective antagonists are acting on peripheral M_2_ or M_3_ receptors. While PD is more classically associated with upper motor neurons, it is possible that antagonism of M_2_ or M_3_ in the periphery somehow blocks or attenuates the expression of tremor at the level of the muscle or lower motor neurons. Further exploration of this possibility of anti-mAChR action at peripheral sites in relevant animal models of disease will be necessary.

While surprising that M_1,_ M_4,_ or M_5_ do not modulate tremor as has been implicated with other motor symptoms of PD, these data implicating unique roles for M_2_ or M_3_ are critical for our understanding of the mAChR mediated mechanisms in PD. These studies help to inform future clinical trials about the potential efficacy of subtype selective mAChRs as they progress to the clinic. Additionally, these studies illustrate the need to mechanistically understand how tremor is induced for the rational design of novel therapeutic strategies for tremor.

## Acknowledgements

We would like to thank Drs. P Jeffrey Conn and Craig Lindsley for the gifts of the experimental compounds used in these experiments and their time in discussing these data.

## Supplemental Information

**Figure S1. Dose Response Curves of Tetrabenazine and Galantamine to Induce Tremulous Jaw Movements. A)** Dose response curve of tetrabenazine (0-10 mg/kg i.p..) to induce tremulous jaw movements. 3 mg/kg produced a threshold induction of TJMs while 10 mg/kg induced a maximal amount. **B)** Dose response curve of galantamine (0-30 mg/kg s.c.) to induce tremulous jaw movements. 1 mg/kg produced a threshold induction of TJMs while 3 mg/kg induced a maximal amount. N=10-15 per group. Data were analyzed with a one-way ANOVA with Dunnett’s multiple comparisons test. *p<0.05, **p<0.05 and ****p<0.0001

**Figure S2. Scopolamine is efficacious in removing Tremulous Jaw Movements. A)** Scopolamine (1 mg/kg i.p..) reduces tetrabenazine induced tremulous jaw movements. **B)** Scopolamine (1 mg/kg i.p..) reduces galantamine induced tremulous jaw movements. N=19-24 per group. Data were analyzed with a paired t-test. ****p<0.0001

